# Prophages integrating into prophages: a mechanism to accumulate type III secretion effector genes and duplicate Shiga toxin-encoding prophages in *Escherichia coli*

**DOI:** 10.1101/2020.11.04.367953

**Authors:** Keiji Nakamura, Yoshitoshi Ogura, Yasuhiro Gotoh, Tetsuya Hayashi

**Affiliations:** Department of Bacteriology, Graduate School of Medical Sciences, Kyushu University, Fukuoka, Japan, 812-8582; Division of Microbiology, Department of Infectious Medicine, Kurume University School of Medicine, Fukuoka, Japan, 830-0011

## Abstract

Bacteriophages (or phages) play major roles in the evolution of bacterial pathogens via horizontal gene transfer. Multiple phages are often integrated in a host chromosome as prophages, not only carrying various novel virulence-related genetic determinants into host bacteria but also providing various possibilities for prophage-prophage interactions in bacterial cells. In particular, *Escherichia coli* strains such as Shiga toxin (Stx)-producing *E. coli* (STEC) and enteropathogenic *E. coli* (EPEC) strains have acquired more than 10 PPs (up to 21 PPs), many of which encode type III secretion system (T3SS) effector gene clusters. In these strains, some prophages are present at a single locus in tandem, which is usually interpreted as the integration of phages that use the same attachment (*att*) sequence. Here, we present prophages integrating into T3SS effector gene cluster-associated loci in prophages, which are widely distributed in STEC and EPEC. Some of the prophages integrated into prophages are Stx-encoding prophages and have induced the duplication of Stx-encoding phages in a single cell. The identified *att* sequences in prophage genomes are apparently derived from host chromosomes. In addition, two or three different *att* sequences are present in some prophages, which results in the generation of prophage clusters in various complex configurations. These “prophages-in-prophages” represent a medically and biologically important type of inter-phage interaction that promotes the accumulation of T3SS effector genes in STEC and EPEC, the duplication of Stx-encoding prophages in STEC, and the conversion of EPEC to STEC and that may be distributed in other types of *E. coli* strains as well as other prophage-rich bacterial species.

**Author summary:** Multiple prophages are often integrated in a bacterial host chromosome and some are present at a single locus in tandem. The most striking examples are Shiga toxin (Stx)-producing and enteropathogenic *Escherichia coli* (STEC and EPEC) strains, which usually contain more than 10 prophages (up to 21). Many of them encode a cluster of type III secretion system (T3SS) effector genes, contributing the acquisition of a large number of effectors (>30) by STEC and EPEC. Here, we describe prophages integrating into T3SS effector gene cluster-associated loci in prophages, which are widely distributed in STEC and EPEC. Two or three different attachment sequences derived from host chromosomes are present in some prophages, generating prophage clusters in various complex configurations. Of note, some of such prophages-in-prophages are Stx-encoding prophages and have induced the duplication of Stx-encoding prophages. Thus, these “prophages-in-prophages” represent an important inter-phage interaction as they can promote not only the accumulation of T3SS effectors in STEC and EPEC but also the duplication of Stx-encoding prophages and the conversion of EPEC to STEC.

## Introduction

Horizontal gene transfer (HGT) is an important mechanism for generating genetic and phenotypic variations in bacteria [1–3]. Phages are major players in HGT, and many temperate phages that confer virulence potential to host bacteria through the transfer of virulence-related genes have been identified [4]. Most temperate phages integrate their genomes into host chromosomes by site-specific recombination to become a part of the chromosomes as prophages (PPs) and enter a lysogenic cycle. Recombination takes place between the homologous sequences of phage and host DNA (*attP* and *attB*, respectively) and is mediated by a phage-encoded integrase [5]. Many bacterial species/strains contain multiple PPs [6–8], providing various possibilities for PP-PP interactions [9, 10]. In particular, *Escherichia coli* strains such as Shiga toxin (Stx)-producing *E. coli* (STEC) strains have acquired more than 10 PPs (up to 21 PPs) [11–14], and some of the PPs are located at the same loci in tandem.

STEC strains cause diarrhea and severe illnesses, such as hemorrhagic colitis (HC) and life-threatening hemolytic-uremic syndrome (HUS). Their key virulence factor is Stx. While there are two subtypes (Stx1 and Stx2) with several variants and STEC produces one or more Stx subtypes/variants [15–18], the known *stx* genes are all encoded by PP genomes. In addition, typical STEC strains share the locus of enterocyte effacement (LEE) locus-encoding T3SS with enteropathogenic *E. coli* (EPEC), and more than 30 effectors have been carried into STEC and EPEC by multiple PPs [19–22]. Thus, EPEC strains are generally regarded as progenitors of typical STEC strains. For example, O157:H7 STEC evolved from an ancestral EPEC O55:H7 through the phage-mediated acquisition of *stx* along with a serotype change [23, 24].

In this study, we initially analyzed the duplicated Stx2-encoding PPs (referred to as Stx-PPs) in STEC O145:H28, one of the major types of non-O157 STEC [25, 26], and found that one of them is integrated into another PP. We then identified its *att* sequence. By subsequent analyses of PPs carrying similar *att* sequences, we show that PP integration in PP (referred to as PP-in-PP) is a genetic event widely occurring in STEC and EPEC and represents a mechanism underlying the evolution and diversification of these bacteria.

## Results

### Integration of inducible and packageable Stx2a phages into a PP integrated into the *ompW* locus in STEC O145:H28

We previously identified 18 PPs in the finished genome of O145:H28 strain 112648 [27], including two Stx2a-PPs found at the *ompW* (P09) and *yecE* loci (P12). The two Stx2a-PP genomes were identical in sequence; thus, they were considered duplicated PPs. As a lambda-like PP (P08) was also found at *ompW*, we initially thought that P08 and P09 had been integrated in tandem. However, by analyzing the potential *att* sequences of the three PPs, we found that while P08 and P12 were integrated into the *ompW* and *yecE* genes with *attL*/*R* sequences of 121 bp and 21 bp, respectively, P09 was integrated into the P08 genome with a 21-bp *attL*/*R* sequence similar to that of P12 (Fig 1a). By analyzing the PPs at the *ompW* and *yecE* loci in O145:H28 strains, we identified another strain (12E129) that carries the same set of PPs: a lambda-like PP at *ompW*, an Stx2a-PP in the PP at *ompW*, and another Stx2a-PP at *yecE* (Fig 1b). The potential *att* sequences of the three PPs were identical to those of the corresponding PPs in strain 112648 (S1 Fig). The genomes of the two Stx2a PPs in strain 12E129 were also nearly identical, excepting the left end. Hereafter, PPs integrated into the same locus are collectively referred to as PPxxx (where xxx denotes the integration locus), such as PP*ompW.*

**Fig 1.**
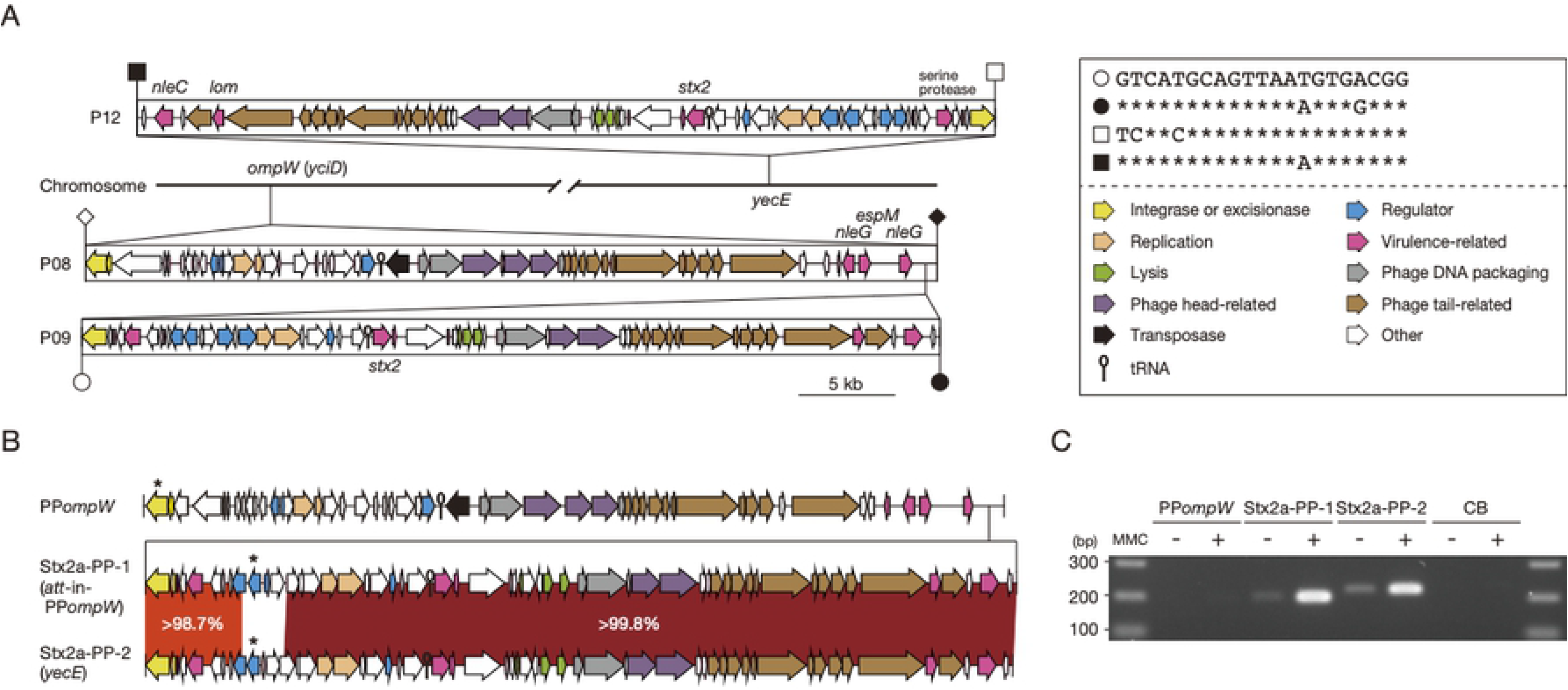
Integration sites of the inducible and packageable duplicated Stx2a phages in two STEC O145:H28 strains. (a) The duplicated Stx2a phages and their *att* sequences in strain 112648. The genome structures of three PPs (P08, P09 and P12) are drawn to scale. The *att* sites of each PP are indicated by open (*attL*) or filled (*attR*) symbols (P08, rhombus; P09, circle; P12, square). The *att* sequences of the Stx2a phages (P09 and P12) are shown in the inset. (b) The genome structures of two Stx2a-PPs and a PP integrated into *ompW* (PP*ompW*) in strain 12E129. Sequence homology between the two Stx2a-PPs is also shown, with their integration sites indicated in parentheses. Homologous regions are indicated by shading with different colors according to sequence identity. The genes that were targeted by the PCR primers used in Fig 1c are indicated by asterisks. (c) Detection of packaged DNA of the three PPs in the DNase-treated lysates of strain 12E129 with (+) or without (−) MMC treatment. The chromosome backbone (CB) region was amplified as a negative control.

To precisely determine the *att* sequences of each PP, we amplified and sequenced the *attP*-flanking regions of excised and circularized genomes of these PPs. Although the two Stx2a-PPs in strain 112648 were indistinguishable, those of strain 12E129 were distinguishable, allowing sequence determination of the *attP*-flanking regions of three PPs from mitomycin C (MMC)-treated cell lysates. This analysis confirmed that the predicted *att* sequences exactly represented those of the three PPs and revealed that these PPs were induced to generate excised and circularized phage genomes by MMC treatment (S1 Fig). The *att* sequences of P08 and P09/P12 in strain 112648 were also confirmed using the same strategy. These results indicate that, in both strains, one of the duplicated Stx2a-PPs has been integrated into PP*ompW*.

We further examined the packageability of these PP genomes into phage particles by PCR analysis of DNase-treated culture supernatants of strain 12E129 with or without MMC treatment (Fig 1c). This analysis detected DNase-resistant genomic DNA of the two Stx2a-PPs, but did not that of PP*ompW*, indicating that the duplicated Stx2a-PPs were both packaged into the phage particles. In a similar analysis of strain 112648, the packaged genome of Stx2a-PP (P09 and/or P12) was detected. That of P08 (PP*ompW*) was also not detected (data not shown), but the reason is currently unknown.

### Dynamics of PP*ompW*s, PPs integrated into PP*ompW*, and PP*yecE*s in STEC O145:H28

To investigate the distribution of PP*ompW*s and the *att* sequences found in two PP*ompW*s (referred to as *att*-in-PP*ompW*) among O145:H28 strains, we selected 64 genomes from 239 strains analyzed in our previous study [27]. This set comprised 8 finished and 56 draft genomes and encompassed seven of the eight clades previously identified in the major lineage (sequence type (ST) 32) and a minor lineage (ST137/6130) of O145:H28, thus largely representing the overall phylogeny of O145:H28 as shown by a whole genome-based maximum likelihood (ML) tree (Fig 2).

**Fig 2.**
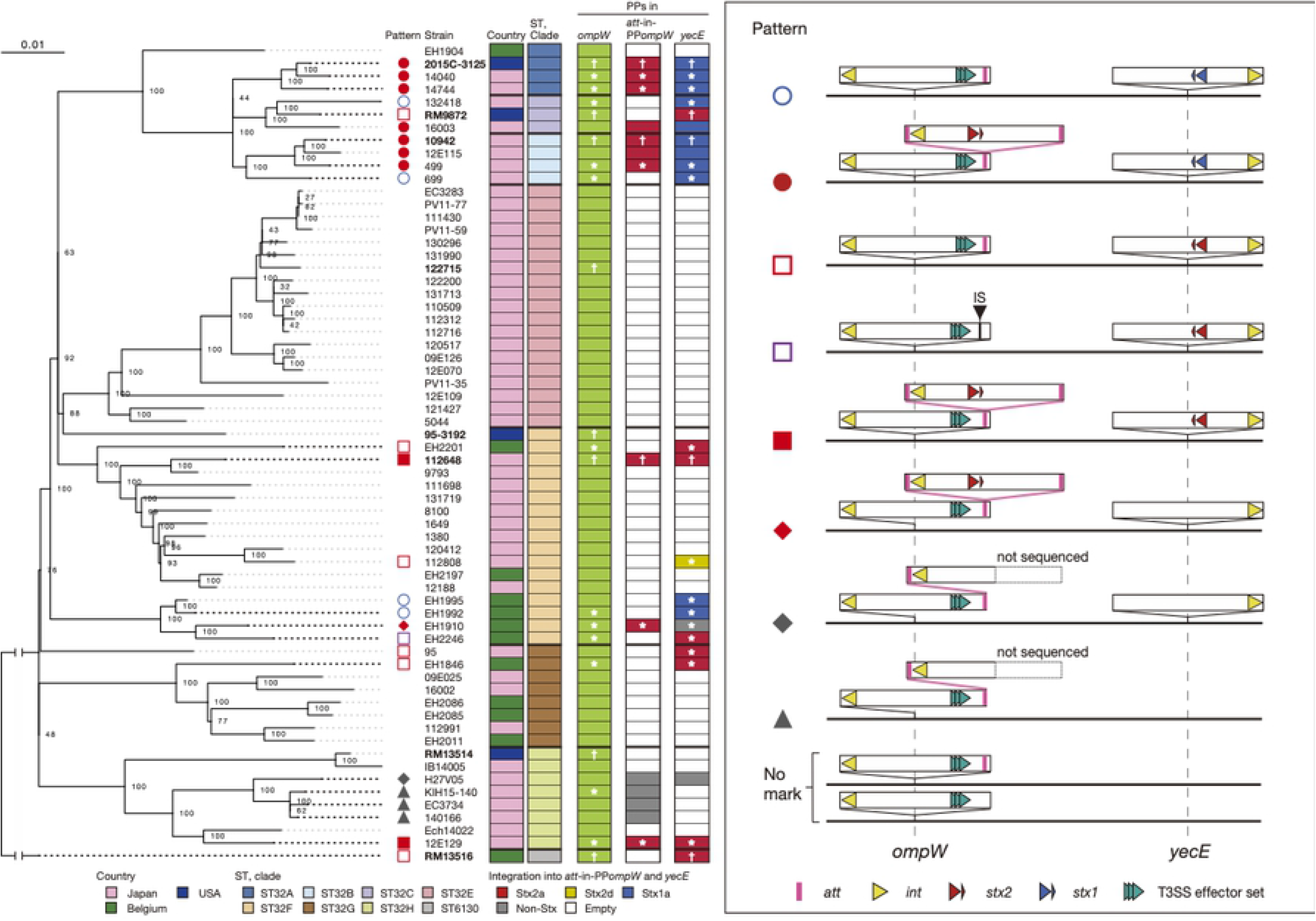
Variation in the PP content at the *ompW*, *att*-in-PP*ompW* and *yecE* loci in STEC O145:H28. In the left panel, an ML tree of 64 O145:H28 strains is shown. Completely sequenced strains are indicated in bold (plasmids were not finished for strain 2015C-3125). The tree was constructed based on the recombination-free SNPs (3,277 sites) identified on the conserved chromosome backbone (CB) (3,961,936 bp in total length) by RAxML using the GTR gamma substitution model [43]. The reliabilities of the tree’ s internal branches were assessed using bootstrapping with 1,000 pseudoreplicates. Along with the tree, the geographic and ST/clade information of strains, the presence or absence of PPs at three loci (*ompW*, *att*-in-PP*ompW* and *yecE*) and the types of PPs at the *att*-in-PP*ompW* and *yecE* loci are shown. PPs sequenced in this study and those in the finished genomes are indicated by asterisks and daggers, respectively. Note that the *att*-in-PP*ompW* sequence is missing from the PP*ompW*s of strains EH2246 and 12E109; a deletion in the latter stain was detected in its draft genome assembly. The bar indicates the mean number of nucleotide substitutions per site. In the right panel, the patterns of the PP content at the three loci are schematically presented. Strains showing each pattern are also indicated in the left panel by diagrams. Note that we detected recombination between the Stx2a-PP at *att*-in-PP*ompW* and a PP present at the *ydfJ* locus that induced a large chromosome inversion in three strains (10942, 499, and EH1910).

PP*ompW*s were present in all 64 strains analyzed, including the two aforementioned strains. All-to-all sequence comparison of the PP*ompW*s from eight finished genomes and 12 PP*ompW*s sequenced in this study revealed that the PP*ompW* genomes were highly conserved, although sequence diversification and segment replacement, probably by recombination, were detected in some parts of several PP*ompW*s (S2a Fig). Further analysis of the 20 PP*ompW*s revealed that all contained the 21-bp *att*-in-PP*ompW* sequence (Fig 2), with one exception where the *att*-containing region had been replaced by an insertion sequence (IS). These results indicate that a PP*ompW* containing *att*-in-PP*ompW* was acquired by an ancestral strain and has been stably maintained in O145:H28.

Examination of PP integration into the *att*-in-PP*ompW* and *yecE* loci in the 64 strains revealed that PPs are integrated into the two loci in 14 and 21 strains, respectively, with marked variation in the PP content between strains (Fig 2). At the *att*-in-PP*ompW* locus, Stx2a-PPs were present in 10 strains and non-Stx-PPs in four strains (all belonging to ST32 clade H). More variable PPs were found at *yecE*: Stx1a-PPs in 11 strains, Stx2a-PPs in eight strains, an Stx2d-PP in one strain, and non-Stx-PPs in two strains. Two aforementioned strains (112648 and 12E129) carrying two duplicated Stx2a-PPs belonged to different ST32 clades, indicating that duplication occurred independently.

All-to-all sequence comparison of 27 PP genomes integrated into the *att*-in-PP*ompW* (n=8; all were Stx2a-PPs) or *yecE* (n=19; 9 Stx1-PPs, 8 Stx2a-PPs, one Stx2d-PP, and one non-Stx-PP) locus revealed that the Stx1a-PP genomes were relatively well conserved, while regions with 2-3% sequence divergence were probably introduced by recombination (S2b Fig). In contrast, the Stx2a-PP genomes were highly variable except for those in the ST32 clade A/B/C strains. Interestingly, although the Stx2a-PP of strain RM9872 (clade C) was integrated into *yecE*, this PP was similar to the Stx2a-PP at *att*-in-PP*ompW* in clade A/B strains. Considering the high conservation of Stx1a-PPs at the *yecE* locus in these clades, it is likely that the Stx1a-PP at *yecE* has been replaced by the Stx2a-PP originally integrated into *att*-in-PP*ompW* in strain RM9872.

### Wide distribution of PP*ompW*s and *att*-in-PP*ompW* in *E. coli*

We next examined the distribution of PP*ompW*s and the *att*-in-PP*ompW* sequence (or sequences similar to it) in the entire *E. coli* lineage by searching for them in 767 publicly available complete *E. coli* genomes. PP*ompW* was found in 44% of the *E. coli* strains examined (338 strains of 92 serotypes; all but O145:H28 and O26:H11 comprised a single ST). Phylogenetic analysis of the *E. coli* strains representing each of the 92 serotypes showed that PP*ompW*s are widely distributed in *E. coli* (Fig 3a). In contrast, after filtering the *att* sequence in *yecE*, 21-bp sequences identical to the *att*-in-PP*ompW* sequence or with a 1-base mismatch (hereafter, collectively referred to as 21-bp sequences) were detected in 150 strains of 20 serotypes belonging to five different *E. coli* phylogroups (Fig 3a and S1 Table). In 145 of the 150 strains, the 21-bp sequence was present in PP*ompW*s. In 28 of the 145 strains (all were serotype O157:H7), two 21-bp sequences were found in two PPs located in tandem at *ompW* (4 strains) or in a PP*ompW* and a PP cluster present at the *mlrA* or *ydfJ* locus (1 and 23 strains, respectively). One atypical O157:H7 strain (PV15-279) carried a PP*ompW*, but the 21-bp sequence was found in the PP cluster at *ydfJ*. The remaining four strains (all were serotype O177:H25) contained no PP*ompW*s, and their 21-bp sequences were found in a PP or a PP cluster at *ydfJ*.

**Fig 3.**
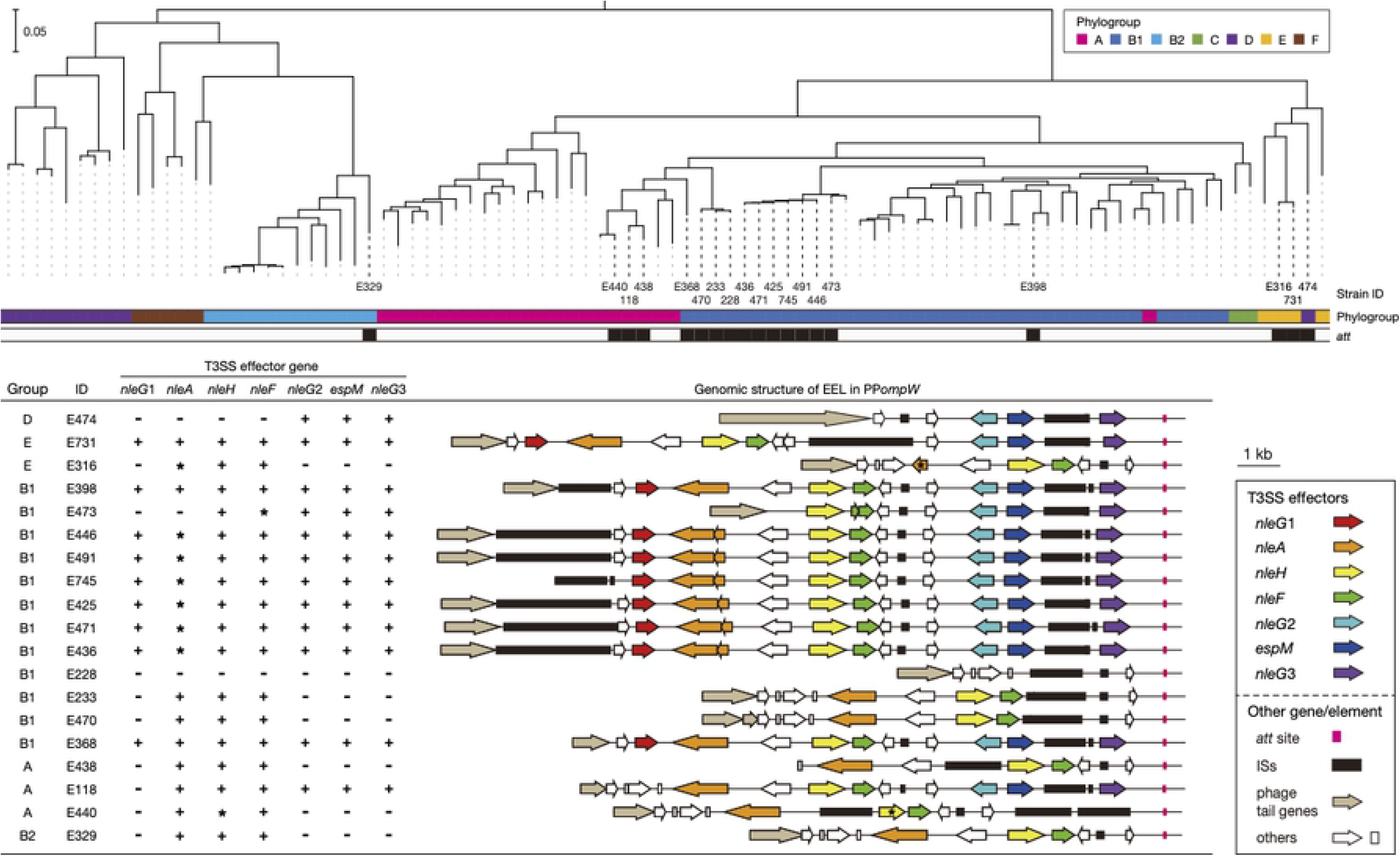
Phylogenetic positions of *E. coli* strains carrying PP*ompW* and the genome structures of their EELs associated with *att*-in-PP *ompW*. In the upper panel, an ML tree of 92 complete genomes of *E. coli* strains that carry PP*ompW* is shown. The tree was constructed based on 109,927 SNP sites in 2,642 core genes and rooted by cryptic Escherichia clade I strain TW15838 (No. AEKA01000000) used as an outlier. Along with the tree, strain IDs used in this paper (see Dataset S2 for more details), phylogroups, and the presence (colored) or absence (open) of *att*-in-PP*ompW* in each strain are indicated. The bar indicates the mean number of nucleotide substitutions per site. In the lower panel, the repertoires of T3SS effector genes that were encoded by the effector exchangeable loci (EELs) in the PP*ompW*s containing *att*-in-PP*ompW* are shown. The genomic structures of EELs are drawn to scale. All effector genes were aligned using BLASTN, and orthologous genes (sequence identity; >90%, coverage; >90%) are indicated by the same color. Genes with over 90% identity but less than 90% coverage and those containing indels and nonsense mutations in the sequence alignment to intact genes are indicated by asterisks.

By examining the 145 strains containing the 21-bp *att*-in-PP*ompW* sequence, we identified additional strains carrying PPs integrated in PP*ompW*s in non-O145:H28 lineages: one O157:H7 strain and two O145:H25 strains (S2 Table). Moreover, as in the two aforementioned O145:H28 strains, the duplication of Stx2-PP and integration of the copies into the *att*-in-PP*ompW* and *yecE* loci occurred in two of the three strains (Stx2d-PP in O157:H7 strain 28RC1 and Stx2a-PP in O145:H25 strain CFSAN004176; S2 Table and S3 Fig), although one of the duplicated Stx2a-PPs in the O145:H25 strain contained a large genomic deletion and its *stx2A* gene was inactivated by multiple insertions and deletions in the coding sequence [14].

### Close association of the *att*-in-PP*ompW* sequence with the PP regions encoding T3SS effector genes

Comparison of the PP*ompW* genomes containing the *att*-in-PP*ompW* sequence (S4 Fig) revealed that while the early regions were relatively well conserved, the late regions were highly variable between PPs due to sequence diversification, deletions and IS insertions. In particular, the PP*ompW* genomes of phylogroup A strains have been highly degraded by deletions. However, multiple T3SS effector genes are present just upstream of the *att*-in-PP*ompW* sequence in all PP*ompW*s except for that in an O182:H25 strain, from which effector genes have apparently been deleted (S4 Fig). Thus, the *att*-in-PP*ompW* sequence is closely linked to the T3SS effector-encoding locus located at the very end of PP*ompW* genomes. Such regions of lambda-like phages encoding various T3SS effector genes are called exchangeable effector loci or EELs [20]. The PP*ompW*s containing the *att*-in-PP*ompW* sequence were also apparently lambda-like phages.

By analyzing T3SS effector genes in the EELs in the 19 PP*ompW* genomes, we identified seven effector genes belonging to the *nleA*, *nleH*, *nleF*, and *espM* families and three *nleG* subfamilies (*G*1-3) (Fig 3). Although there were variations in the effector gene repertoire between PP*ompW*s and gene inactivation due to various types of mutations (mostly deletions) was detected in several PP*ompW*s, a similar set of effector genes was found at the PP*ompW* EELs. As one or more IS elements were present at all EELs, the variation in effector gene repertoire was probably generated by IS insertion-associated events. The conservation patterns of effector genes among the 19 PP*ompW* genomes suggest that the EELs of O157:H7 strain Sakai (phylogroup E; E731 in Fig 3) and EPEC O76:H7 strain FORC_042 (phylogroup B1, E398 in Fig 3) represent the ancestral structure encoding seven effector genes.

It should be noted that all strains carrying a PP(s) that contained the 21-bp *att* sequence and the associated EEL(s) possessed the *eae* gene, a marker gene of the LEE (S1 Table), indicating that they are all EPEC or typical (LEE-positive) STEC.

### PP clusters that contained PPs carrying the *att*-in-PP*ompW* sequence and identification of additional *att* sites in PP genomes

In four of the aforementioned 28 O157:H7 strains that contained two 21-bp sequences identical or nearly identical to *att*-in-PP*ompW*, the sequences were each present in the EEL-associated region of two PP*ompW* genomes integrated in tandem (Fig 4a and S5 Fig). In these strains (as represented by FRIK2069 in Fig 4a), while one of the EELs encoded an effector gene set similar to that of other PP*ompW* EELs, the other encoded an *nleG* variant different from the three *nleG* families at other PP*ompW* EELs.

**Fig 4.**
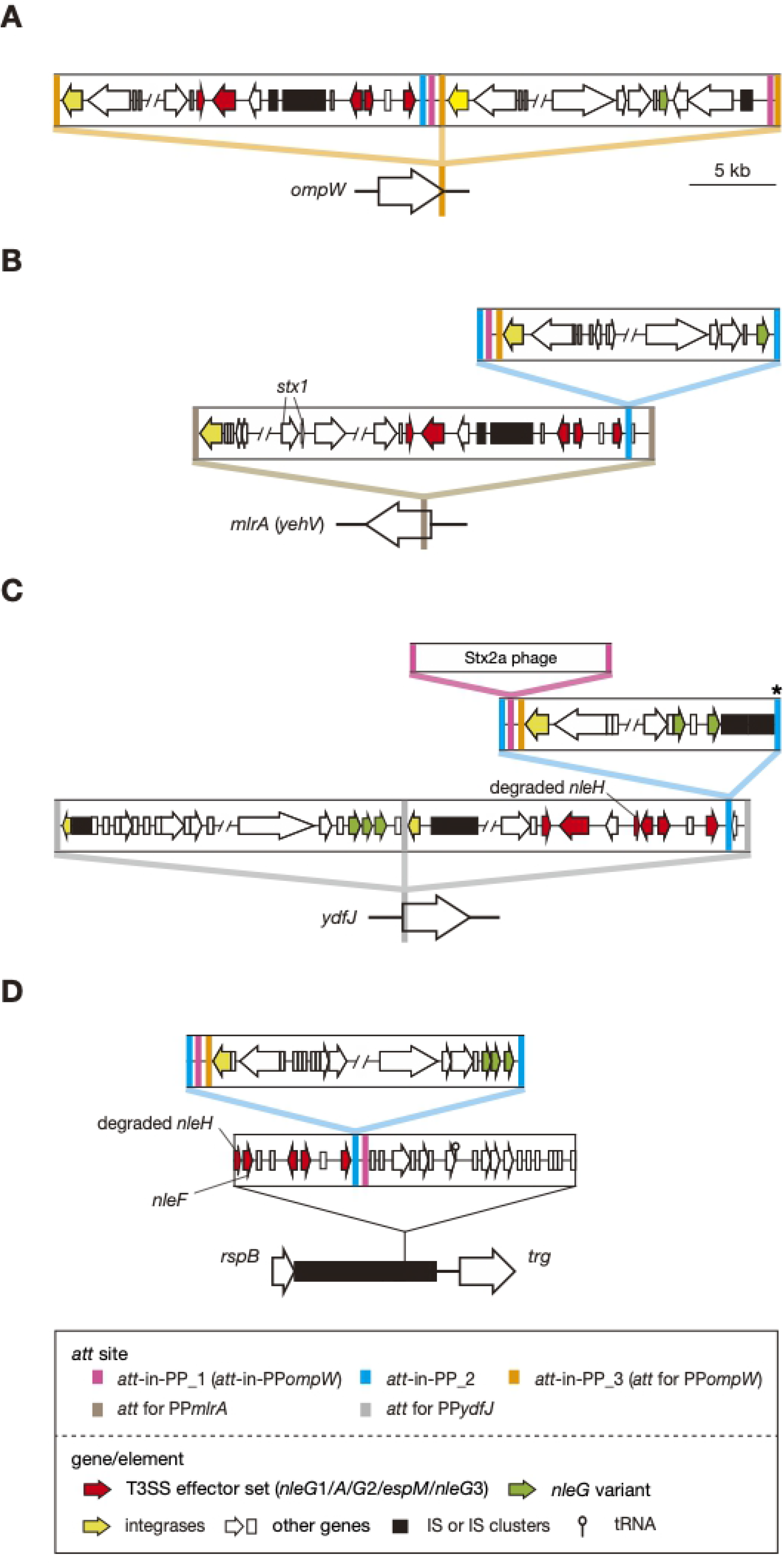
PP clusters that contained PP carrying potential *att* sequences in O157:H7 and O177:H25 strains. The genomic structures of three representative PP clusters of the 33 PP clusters found in O157:H7 and that of O177:H25 strains are shown (A, strain FRIK2069; B, strain FRIK944; C, atypical O157:H7 strain PV15-279; D, O177:H25 strain SMN152S1). The identified *att* sequences, coding sequences (CDSs) (including pseudogenes), and ISs in each PP are indicated. T3SS effector genes found in the PP*ompW* EELs (Fig 3) and other effector genes (*nleG* variants) are distinguished by different colors. In panel C, the *att* sequence indicated by an asterisk is truncated by an IS insertion, and integration of an Stx2a-PP into the *att*-in-PP_1 site is schematically presented. The genome structures of all PP clusters identified in this analysis are illustrated in S5 Fig.

In 24 O157:H7 strains, one 21-bp sequence was present in PP*ompW,* and the other was present in PP-in-PP clusters comprising two to four PPs. In one strain (FRIK944; Fig 4b), the PP cluster was present at *mlrA* (synonyms: *yehV*) and comprised two PPs, an Stx1-PP and a lambda-like PP. By analyzing the *attL*/*R* sites of each PP, we found that while Stx1-PP is integrated into *mlrA* [9], the lambda-like PP is in Stx1-PP, using the 96-bp *att* sequence (referred to as *att*-in-PP_2; see S6 Fig for the sequence) associated with an EEL similar to PP*ompW* EELs. The lambda-like PP also contained an *nleG* variant, but the 21-bp sequence was present between *attL* and the integrase gene and was not associated with the *nleG* variant. As it is now known that the 21-bp sequence is present in a PP genome other than PP*ompW* genomes, we hereafter refer to it as *att*-in-PP_1. Intriguingly, between *att*-in-PP_1 and the integrase gene of the lambda-like PP, the 121-bp *att* sequence for PP*ompW*s was present. Although PP integration into the 121-bp sequence in PP genomes has yet to be identified, this sequence can serve as a potential *att* site in PP genomes. We therefore refer to it as *att*-in-PP_3.

In the remaining 23 strains, PP clusters comprising two to four PPs were present at *ydfJ* (Fig 4c; see S5 Fig for other strains). In these strains, one or two lambda-like PPs, which carry EELs similar to the PP*ompW* EELs or encode multiple *nleG* variants, were integrated into *ydfJ* (see S7 Fig for the *att* sequences). The former type of EEL was associated with *att*-in-PP_2, into which another lambda-like PP was integrated. Similar to the PP integrated into PP*mlrA* (Fig 4b), the PPs integrated into PP*ydfJ* contained the *att*-in-PP_1 and *att*-in-PP_3 sequences downstream of the integrase gene and encoded *nleG* variants at the opposite PP end. Moreover, in one of the 23 strains (PV15-279, an atypical O157:H7 strain [28]), an Stx2a-PP was integrated into the *att*-in-PP_1 of the PP integrated into PP*ydfJ* (Fig 4c).

Among the four aforementioned O177:H25 strains that contained the 21-bp *att*-in-PP_1 sequence, a similar but slightly different pattern of PP integration into PP genomes was observed (Fig 4d). In these strains, the *att*-in-PP_1 sequence was found in a PP-like region that probably represents two highly degraded PP genomes integrated in tandem between the *rspB* and *trg* genes. EELs similar to the PP*ompW* EELs, *att*-in-PP_2 and *att*-in-PP_1 were found in this order, and a lambda-like PP was integrated into *att*-in-PP_2. Moreover, the lambda-like PPs integrated into *att*-in-PP_2 contained the *att*-in-PP_1 and *att*-in-PP_3 sequences and multiple *nleG* variants, as PPs integrated into PP*mlrA* or PP*ydfJ*s (Fig 4d). This finding indicates that the distribution of these three *att* sequences in PP genomes is not limited to O157:H7 strains.

### Origins of *att*-in-PP sequences

Finally, to explore the origins of these *att*-in-PP sequences, we compared their flanking sequences with *E. coli* chromosome sequences. The *att*-in-PP_1-flanking sequences in PP*ompW*s and other PPs (all are integrated into PPs as shown in S5 Fig) were highly conserved, implying that these sequences have a common origin (S8 Fig). Moreover, the 100-bp sequences including the *att*-in-PP_1 sequence showed a notable similarity (87% identity) to the corresponding *yecE* region (Fig 5, see S8 Fig for sequence alignment), suggesting that the *att*-in-PP_1 and its flanking sequence originated from the *yecE* locus.

**Fig 5.**
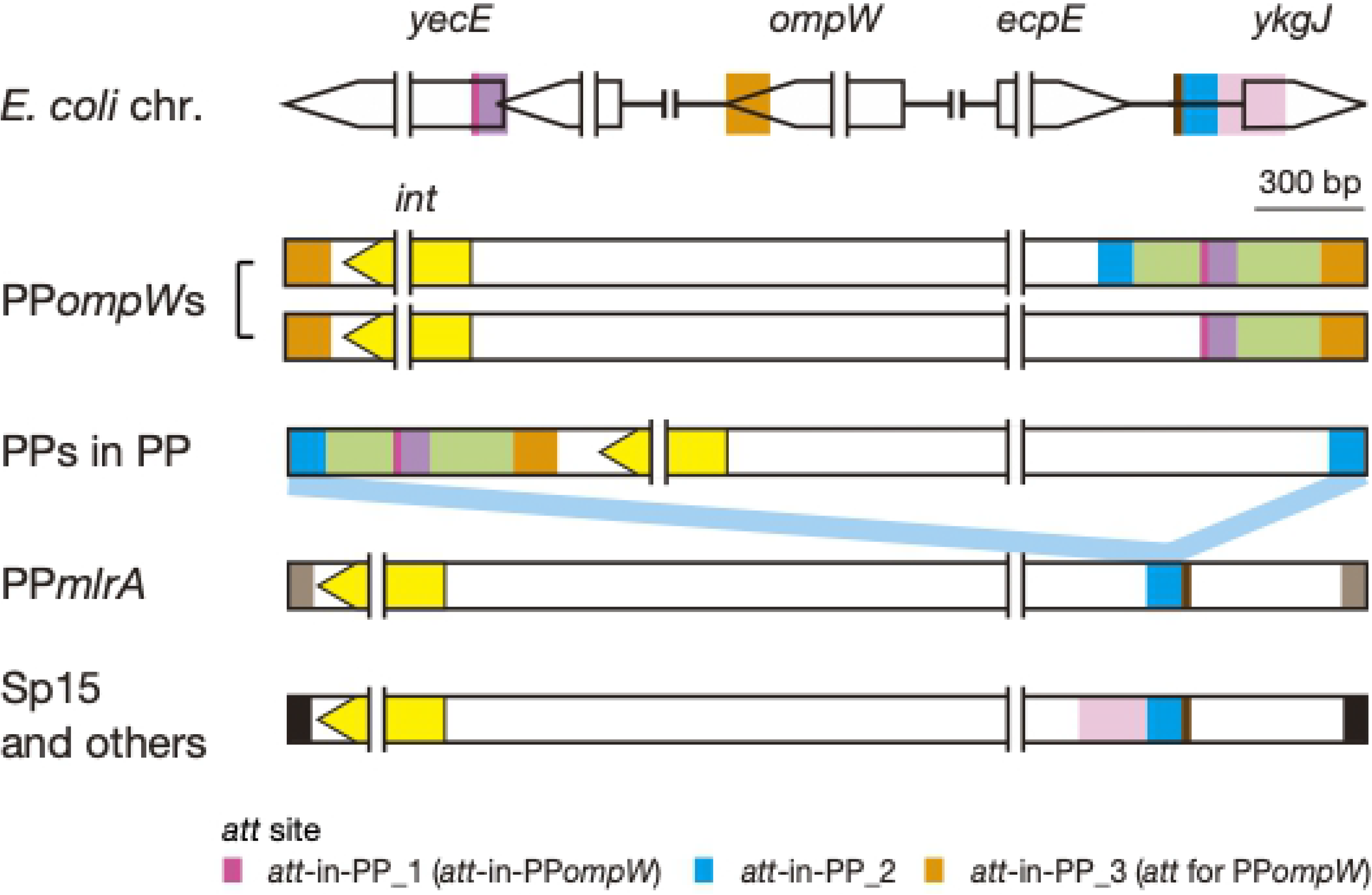
Locations of the *att*-in-PP sequences in PPs and the PP genome regions homologous to *E. coli* chromosome regions. Three loci in the *E. coli* chromosome showing sequence homology to three identified *att*-in-PP sequences and their flanking sequences are shown at the top. The left- and right-end regions of representative PPs that contained the *att*-in-PP sequences are shown below. Homologous sequences are indicated by the same color. The color used for each *att*-in-PP sequence is the same as that used in Fig 4. See Fig 4 for the details of “PPs-in-PPs” and “PP*mlrA*” and S9 Fig for information on “Sp15 and other PPs”. Alignments of the *att*-in-PP_1 and *att*-in-PP_2 sequences and their flanking sequences with corresponding chromosome sequences are shown in S8 and S9 Figs, respectively.

Sequence similarity was also detected between the 96-bp *att*-in-PP_2 sequence in PP*ompW*s and the *ykgJ*/*ecpE* intergenic region of the *E. coli* chromosome (78% identity) (Fig 5). As the homologous sequence extended to 125 bp in PP*mlrA*, we performed an additional search of *E. coli* complete genomes and identified seven *att*-in-PP_2-containing PPs, although this search was limited to six STEC genomes fully annotated for PPs (S9 Fig). The identified PPs included the Stx1a-PP (Sp15) at *mlrA* of O157:H7 strain Sakai [11], the aforementioned duplicated Stx2a-PPs of O145:H28 strain 112648, duplicated Stx2a-PPs of the atypical O157:H7 strain PV15-279 (one in PP*ompW* and the other in *yecE*; carrying a T3SS effector gene), and two PPs in O26:H11 and O111:H8 STEC strains [12] (at *ydfJ* and *ssrA*, respectively; the former carries a T3SS effector gene). In these seven PPs, homologous sequences further extended to 309 bp with 84% identity (Fig 5). Contrary to the observation for *att*-in-PP_1 and its flanking sequences, there was notable diversity in the *att*-in-PP_2 sequence (20/96 polymorphic sites) between the PP*ompW*s, PP*mlrA* in FRIK944, and the other seven PPs (S9 Fig). These findings indicate that the *att*-in-PP_2 and its flanking sequences originated from the *ykgJ*/*ecpE* intergenic region on the chromosomes of *E. coli* or its close relatives, but acquisition of the sequences by phages might have occurred multiple times.

The 121-bp *att*-in-PP_3 sequence was found in many of the PPs-in-PPs identified in this study (Figs 4 and 5, and S5 Fig) and showed 81% identity to the *E. coli ompW*, suggesting that its possible origin is also the chromosome of *E. coli* or its close relatives. Interestingly, PP*ompW*s and many other PPs-in-PPs contained two or three *att*-in-PP sequences in the same order. The sequences between the *att*-in-PP sequences (indicated by green in Fig 5) were also conserved (up to a 5-single nucleotide polymorphism (SNP) difference); however, the location of the *att*-in-PP set in PP*ompW*s was different from that of other PPs integrated in PPs. This finding suggests that the region encompassing three (or two) *att*-in-PP sequences was once acquired by either type of phage and spread to the other by recombination or some other mechanisms.

## Discussion

As summarized in Fig 6, we identified various PP integration patterns in STEC and EPEC strains, including PP integration into PPs. Most temperate phages are integrated into host genomes by integrase-mediated recombination between *attP* and *attB.* Tandem PP integration can occur if the two phages share the same *attB* site. In contrast to this traditional view of the mechanism for generating tandem PPs, this study identified many PPs that contain *att* sequences, which allow another PP to be integrated into their genomes, forming a PP-in-PP configuration. The combination of the two integration mechanisms generates more complex PP clusters in host genomes (combination of tandem PPs and PPs-in-PPs). Frequent colocalization of multiple *att*-in-PP sequences potentially generates much more variation than detected in this study. These *att*-in-PP sequences originated from the host chromosome, providing more opportunities for lysogenization to incoming phages and allowing the duplication of PPs encoding medically or biologically important genes, such as *stx*. There may be some previously unrecognized interaction(s) between integrating PPs and their “host” PPs. Analyses of such interactions as well as the mechanisms of incorporating *att*-in-PP sequences from host chromosomes are worthy of future studies to better understand the processes of PP-in-PP formation.

**Fig 6.**
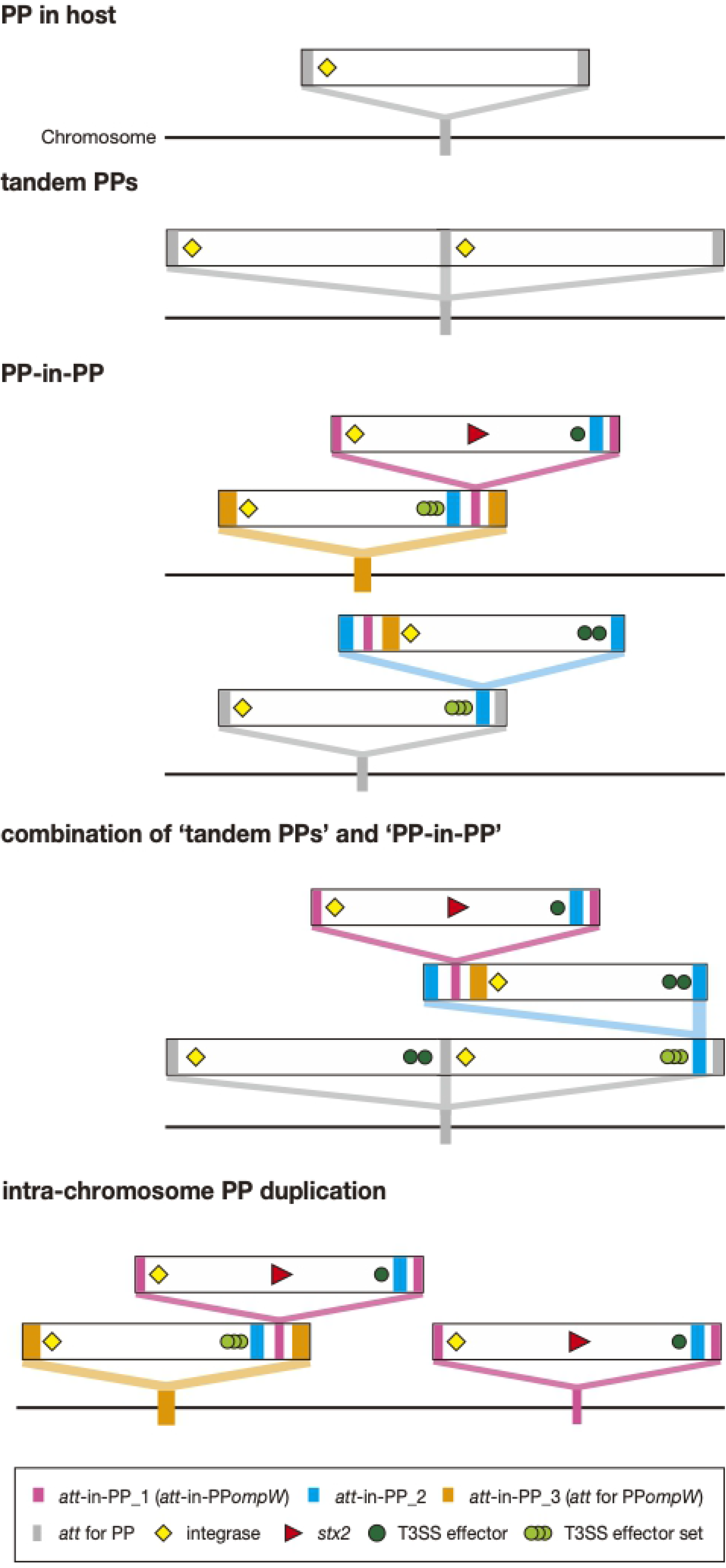
Summary of the variable PP integration patterns found in this study.

Notably, most *att*-in-PP sequences identified are linked to EELs that encode multiple effector genes for the LEE-encoded T3SS, and PPs integrated into *att*-in-PPs often carry effector genes. Thus, the PP-in-PP system has promoted the accumulation of effector genes in EPEC and STEC strains [21, 29] and can promote further accumulation of these genes, which may increase the pathogenicity of these strains [22, 30]. Furthermore, a significant portion of the PPs integrated in *att*-in-PP_1 (13/18) encoded *stx* genes, indicating that the *att*-in-PP_1 sequence has promoted the acquisition of *stx* genes and thus the conversion of EPEC to typical STEC, even if the *yecE* locus, the origin of *att*-in-PP_1 and one of the integration hot spots of Stx-PP [12, 31–33], has been occupied by another PP.

In conclusion, the findings obtained here highlight that PP integration systems are much more complicated than previously recognized and provide additional insights into the evolution of EPEC and STEC and their pathogenicity. It is also possible to find similar PP integration patterns in other types of *E. coli* and other PP-rich species if PP clusters are carefully investigated. Similar integration systems could also be found for genetic elements utilizing integrase-mediated integration mechanisms, such as integrative and conjugative elements (ICEs) [34].

## Material and Methods

### Bacterial strains

The 64 O145:H28 strains analyzed in this study are listed in S3 Table. Of these, 59 were from our laboratory stock, which were genome-sequenced in our previous study [27], and 5 were completely genome-sequenced stains (the plasmid genome was not finished in strain 2015C-3125), the genome sequences of which were downloaded from the NCBI database. To construct the completely genome-sequenced *E. coli* strain set, a total of 875 complete genomes were downloaded from the database (accessed on the 20th of July 2019). After excluding laboratory, commercial and re-sequenced strains and substrains, the 767 strains listed in S4 Table were used for analysis. Annotation was carried out using the DDBJ Fast Annotation and Submission Tool (DFAST) [35], if necessary.

### Extraction of total cellular and phage DNA

Bacterial cells were grown overnight to the stationary phase at 37°C in lysogeny broth (LB) medium. For prophage induction, cells were grown to the late log phase (0.7-0.9 OD_600_), and MMC was added to the culture to a final concentration of 1 μg/ml. After a 3-hr incubation, aliquots of the culture were isolated, and the cells were collected by centrifugation. Total cellular DNA was extracted from the cells using the alkaline-boiling method and used for PCR analyses. Phage particles were isolated from the culture supernatant after a 3-hr incubation with MMC. The culture was first treated with chloroform, and bacterial cell debris was removed by centrifugation. The supernatant was filtered through a 0.2-μm-pore-size filter (Millipore) and incubated with DNase I (final concentration: 400 U/ml, TaKaRa) and RNase A (50 μg/ml, Sigma) at 37°C for 1 hr. After inactivating DNase I by incubation at 75°C for 10 min and adding EDTA (5 mM, Nacalai Tesque), the sample was treated with proteinase K (100 μg/ml; Wako) and used as packaged phage DNA. Total cellular DNA and packaged phage DNA from MMC-untreated cultures were prepared with the same protocol. The primers used in these analyses are listed in S5 Table.

### Analyses of PP integration and sequencing of PP genomes

PP integration into the *ompW*, *att*-in-PP*ompW* (later renamed *att*-inPP_1) and *yecE* loci in 56 O145:H28 draft genomes was first examined by a BLASTN search as outlined in S10a Fig. The integration of Stx PPs into *att*-in-PP*ompW* and/or *yecE* was determined by long PCR amplification using primers targeting the *stx* genes and sequences adjacent to these integration sites, as schematically shown in S10b Fig. The products of long PCR were used for sequence determination of each PP. The primers used in this analysis are listed in S6 Table.

Sequencing libraries were prepared for each product of long PCR (ranging from 15 to 33 kb) using the Nextera XT DNA Sample Preparation Kit (Illumina) and sequenced on the Illumina MiSeq platform to generate paired-end (PE) reads (300 bp x 2). PP genomic sequences were obtained by assembling and scaffolding Illumina PE reads using the Platanus_B assembler (v1.1.0) (http://platanus.bio.titech.ac.jp/platanus-b) [36]; then, gaps were closed by Sanger sequencing PCR products that spanned the gaps. Annotation of all PP genomes was carried out with DFAST, followed by manual curation using IMC-GE software (In Silico Biology). All sequences have been deposited in the DDBJ/EMBL/GenBank databases under the accession numbers listed in S3 Table. GenomeMatcher (v2.3) [37] was used for genome sequence comparison and to display the results.

### Searches for PP*ompW*s and *att*-in-PP*ompW* sequences in the complete *E. coli* genomes

Serotypes and *eae* subtypes of the 767 complete *E. coli* strains were determined by BLASTN as previously described [29]. Systematic ST determination was performed by a read mapping-based strategy using the SRST2 program [38] with default parameters. Read sequences of the complete genomes were simulated with the ART program (ART_Illumina, version 2.5.8) [39]. The genomes whose ST was not precisely defined (possible ST containing a novel allele, an uncertain ST, and no STs in the present database) were reanalyzed using MLST 2.0 with “Escherichia coli #1” schemes [40] (https://cge.cbs.dtu.dk/services/MLST/).

The presence of PP*ompW* and the *att*-in-PP*ompW* sequence was examined in the complete genomes by a BLASTN-based search as follows. The presence of PP*ompW* was determined using two query sequences: one was the integrase gene of O145:H28 strain 112648 (EC112648_1574) (thresholds: >90% identity and >90% coverage), and the other was the *ompW*-containing region on the chromosome of *E. coli* K-12 (No. NC_000913; nucleotide positions 1,314,020-1,315,224; no PP integration) to examine the absence of PP insertion into the *ompW* locus (threshold: >85% identity and <60% coverage). When either the integrase gene or the *ompW* locus split by some insertion was detected, we analyzed the gene organization of these regions to determine if PP*ompW* was present. The search for the 21-bp *att*-in-PP*ompW* sequence (5’-GTCATGCAGTTAAAGTGGCGG-3’) (S1c Fig) was performed with the blastn-short task option (thresholds: >95% identity and 100% coverage). The 21-bp sequences in the *yecE* gene, which were similar to the *att*-in-PP*ompW* sequence, were removed.

### SNP detection and phylogenetic analysis

The SNP sites (3,277 sites) of the core genomic sequences of the 64 O145:H28 strains were detected by MUMmer [41], followed by filtering recombinogenic SNPs by Gubbins [42], and used for reconstruction of an ML tree in RAxML [43] with the GTR gamma substitution model as previously described [27]. To reconstruct the phylogeny of the *E. coli* strains carrying PP*ompW*, we used 92 *E. coli* strains representing each of the 92 serotypes that contained PP*ompW*-carrying strains. Strains in which *att*-in-PP*ompW* was detected were preferentially selected from the serotypes that contained multiple strains. *Escherichia* cryptic clade I strain TW15838 (No. AEKA01000000) was used as an outgroup. The core genes (n=2,642) of these strains, which were defined as the genes present in 100% of strains, were identified by Roary [44], and their concatenated sequence alignments were generated by the same software. Based on the alignment (109,927 SNP sites in total), an ML tree was constructed using RAxML as described above. Phylogroups of the strains were determined by ClermonTyping [45]. ML trees were displayed and annotated using iTOL [46] or FigTree (v1.4.3) (http://tree.bio.ed.ac.uk/software/figtree/).

## Acknowledgements

This research was supported by AMED under Grant Number 20fk0108065h0803 to T.H., and a KAKENHI from the Japan Society for the Promotion of Science (18K07116) to K.N. We thank M. Horiguchi, M. Kumagai, Y. Nagayoshi, and K. Ozaki for providing technical assistance. We also thank the EHEC working group in Japan for providing O145:H28 strains.

## Author Contributions

**Conceptualization:** Nakamura K, Hayashi T.

**Data curation:** Nakamura K

**Formal analysis:** Nakamura K, Ogura Y, Gotoh Y.

**Funding acquisition:** Nakamura K, Hayashi T.

**Investigation:** Nakamura K

**Methodology:** Nakamura K, Ogura Y, Gotoh Y.

**Project Administration:** Hayashi T.

**Resources:** Nakamura K, Ogura Y.

**Visualization:** Nakamura K

**Writing - Original Draft Preparation:** Nakamura K

**Writing - Review & Editing:** Ogura Y, Gotoh Y, Hayashi T.

## Supporting information

**S1 Fig. Determination of the *attP* sequences of PPs integrated in *ompW*, PP*ompW*, and *yecE* in O145:H28 strains 12E129 and 112648.** (a) Schematic representation of the PCR strategy used to amplify the *attP*-flanking region (left panel) and the locations of PCR primers used for each PP (right panel). (b) PCR detection of excised and circularized PP genomes. Total cellular DNA isolated from MMC-treated (+) or MMC-untreated (−) cells was analyzed. A chromosome backbone (CB) region was amplified as a positive control. (c) The *att* sequences of the three PPs in strain 12E129. The *attP*-containing sequences obtained by sequencing the PCR products shown in S1b Fig were aligned with the *attR*-, *attL*-, *attB*-containing sequences to define the *att* sequences of each PP. The *ompW* sequences of strain K-12 MG1655 (accession No. NC_000913) and the *yecE* sequence of O145:H28 strain 122715 (accession No. AP019708), in which no PPs were integrated, were used as the *attB* sequences, respectively (indicated by a dagger and a section mark, respectively). The defined *att* sequences are indicated by uppercase letters.

**S2 Fig. All-to-all genome sequence comparison of PP *ompW*s and that of PP*att*-in-PP*ompW*s and PP*yecE*s found in 64 O145:H28 strains.** Dot plot matrixes of the concatenated sequences of the 20 PP*ompW* genomes (a) and 27 PP*att*-in-PP*ompW* and PP*yecE* genomes (b) found in 64 O145:H28 strains are shown. Strain names and information on the ST and ST32 clade of each strain are indicated. Sequence identities are indicated by a heatmap. In panel A, the nucleotide sequences between *att*-in-PP*ompW* and *attR* (approximately 428 bp in length) were excluded from this analysis because the sequences of this region in three strains (499, EH1910, and KIH15-140) were not determined. In panel B, the subtype of Stx encoded by each PP and the integration site of each PP are indicated. PP groups sharing similar genomic sequences are framed by boxes.

**S3 Fig. Variation in the sequences of the Stx2-encoding PP *att*-in-PP*ompW* and PP *yecE* genomes found in *E. coli* strains analyzed in this study.** Dot plot matrixes of the concatenated sequences of the PP*att*-in-PP*ompW* and PP*yecE* genomes encoding Stx2 are shown. The genome sequences of a pair of PP*att*-in-PP*ompW* and PP*yecE* found in two O145:H28 strains, that in an O157:H7 strain and that in an O145:H25 strain were compared. The names of host strains, Stx2 subtypes, and integration site of each PP are indicated. PPs in the same strain are framed by boxes. Sequence identities are indicated by a heatmap.

**S4 Fig. Comparison of the PP *ompW* genomes containing the *att*-in-PP*ompW* sequence.** In the left panel, along with the same ML tree as shown in Fig 3, *E. coli* strain ID, phylogroup (PG), and the presence (colored) or absence (open) of the *att*-in-PP*ompW* sequence in each *E. coli* are indicated. In the right panel, the genome structures of PP*ompW*s containing *att*-in-PP*ompW* are drawn to scale. A large chromosome inversion resulting from recombination between PP*ompW* and another PP in two strains is indicated by an asterisk (E473 and E471), and relevant PP regions are shown. Homologous regions and sequence identities are depicted by shading with a color gradient. The Stx2a-PPs integrated into the *att*-in-PP*ompW* locus in strains E474 and E118 are schematically indicated.

**S5 Fig. Variation in the PP integration patterns in the PP clusters that contained PPs carrying potential *att* sites.** In the left table, a list of 33 strains that possessed PP clusters that contained PPs carrying the 21-bp sequence identical or nearly identical to the *att*-in-PP*ompW* sequence is provided. In the right panel, the patterns of PP integration are schematically illustrated. Strains showing each pattern are also indicated in the left table. CDSs shown by colored triangles include pseudogenes. The 21-bp sequence (renamed *att*-in-PP_1) and other *att* sequences are indicated. Among these sequences, the two indicated by an asterisk are truncated by IS insertion. Several *att* sequences are missing because of deletions. The T3SS effector set (light green triangles) consists of any of the seven effector family/subfamily genes that are encoded by the PP*ompW* EELs shown in Fig 3. More detailed genomic structures of four PP clusters (indicated in bold in the left table) are presented in Fig 4. Types a, c and d include a minor variation; homologous recombination between the second PP*ompW* and the first PP*ydfJ* (type a2), integrase-deficient PP*ydfJ*s with or without additional PP integration in tandem (types c2 and c3, respectively), and a region comprising two degraded PPs integrated in tandem between the *rspB* and *trg* genes without PP integration into the *att*-in-PP_2 locus (type d2) are shown.

**S6 Fig. Variation in the PP integration patterns in the PP clusters that contained PPs carrying potential *att* sites.** (a) Locations of the *att*-in-PP_2 sequences in representative PP genomes. (b) Comparison of the nucleotide sequence of *att*-in-PP_2 among the PPs shown in panel A.

**S7 Fig. Variation in the PP integration patterns in the PP clusters that contained PPs carrying potential *att* sites.** (a) Schematic representation of the *ydfJ*-flanking region and the PP clusters present at the *ydfJ* locus in three *E. coli* strains. Because both integrase genes of the PP*ydfJ*s in strain PV15-279 (PP*ydfJ*-L and PP*ydfJ*-R) have been inactivated by IS insertion, the PP*ydfJ*-L of O26:H11 strain 11368 was used for sequence determination of the *attP*-flanking region of PP*ydfJ* by sequencing a PCR amplicon obtained with two primers (indicated by red and blue arrows). (b) The *att* sequences of the four PP*ydfJ*s. The *attP*-containing sequence of the PP*ydfJ*-L of strain 11368 was aligned with the *attR*-, *attL*-, and *attB*-containing sequences to define the *att* sequences of each PP. The *ydfJ* sequence of O104:H4 strain C227-11, in which no PPs were not integrated, was used as the *attB* sequence. The 18- or 19-bp *att* sequence that we defined is indicated by uppercase letters.

**S8 Fig. The *att*-in-PP_1 and its flanking sequences in PPs and comparison with the *E. coli yecE* sequence.** (a) The locations of the *att*-in-PP_1 (initially called *att*-in-PP*ompW*) sequences in the genomes of six PP*ompW*s and three other PPs integrated in PPs and in the *yecE* locus of *E. coli* O145:H28 strain 122715. The 21-bp *att*-in-PP_1 sequence and the additional 79-bp sequence homologous to the *yecE* gene are indicated by red and purple, respectively. The *att*-in-PP_2 and *att*-in-PP_3 are also indicated by blue and orange, respectively. The sequences of the two regions indicated by green are conserved between PPs with up to 5 SNPs. The lengths of the two regions are 185 bp (left) and 228 bp (right). (b) Alignment of the 100-bp sequences homologous to the *yecE* locus in the nine PPs shown in panel A with the corresponding sequence of the *yecE* locus of strain *E. coli* O145:H28 strain 122715. The 21-bp *att*-in-PP_1 sequence is indicated by uppercase letters. The 100-bp sequences of these PPs were 87% identical to the *yecE* sequence.

**S9 Fig. The *att*-in-PP_2 sequence and its flanking sequences.** (a) The locations of the *att*-in-PP_2 sequences (blue) in seven PP genomes and on the chromosome of *E. coli* K-12 strain MG1655. The 96-bp *att*-in-PP_2 sequences and their flanking sequences (184 bp and 29 bp in length) homologous to the *ykgJ*-*ecpE* region on the *E. coli* MG1655 chromosome are indicated by blue, pink, and dark brown, respectively. The presence of *stx* and T3SS effector genes in each PP is also indicated. (b) Alignment of the att-in-PP_2 and its flanking sequences in the PPs shown in panel a with the corresponding sequence of the *ykgJ*-*ecpE* region on the *E. coli* MG1655 chromosome. Only the PP genomic regions homologous to the *ykgJ*-*ecpE* region are shown. The 184-bp regions (pink) of PPs show 83% sequence identity with the *ykgJ*-*ecpE* region. Note that the 96-bp *att*-in-PP_2 (blue; indicated by uppercase letters) contained 20 SNPs.

**S10 Fig. Procedures used to determine the PP integration into the *ompW*,*att*-in-PP*ompW* (later in the manuscript, renamed *att*-in-PP_1) and*yecE* loci.** (a) Analysis of PP integration by a BLASTN search. Draft genomes of O145:H28 (n=56) were searched by BLASTN, using the sequences of the *attL*- and *attR*-containing regions of the PPs at *ompW*, *att*-in-PP*ompW* and *yecE* in strain 112648 (P08L/R, P09L/R and P12L/R, respectively) as queries. Each query sequence was composed of the sequences from the host chromosome and PP (60 bp each) with the *att* sequence determined in this study (121 bp for P08 and 21 bp for P09/P12) located between them. PP integration at each locus was considered positive when *attL*- and *attR*-containing sequences were both detected (identity threshold: >95%). PP integration in all but two genomes was determined by this analysis. In strains EH1910 and H27V05, although PPs integrated into *yecE* (PP*yecE*) were detected, PP*ompW* was not detected. Unexpectedly, however, the P09L/R sequences (corresponding to the *attL*- and *attR*-containing sequences of the PP-in-PP*ompW*) were detected in EH1910, and a partial P09 *attL* sequence (74.5% coverage) was detected in H27V05. Therefore, the *ompW* and *att*-in-PP*ompW* loci of the two genomes were defined as ‘Others’, and subjected to long PCR analysis along with the identified PPs. (b) Long PCR analysis and sequence determination of PPs. Strategies for five types of analysis are shown. Type I analysis: The genomes of PPompWs that did not contain PPs were divided into three segments and amplified by three long PCRs to obtain the PCR products for genomic sequence determination. Note that the left and right segments included the left and right PP*ompW*-chromosome junctions, respectively (the same strategy was employed in Types II-V analyses). Type II analysis: The genomes of PP*ompW*s that contained an Stx-PP were amplified together with the Stx-PP genomes using 5 or 6 primer pairs to confirm the presence of these PPs and to obtain the PCR products for genome sequence determination. Two primers targeted the *stx* gene (*stx1* or *stx2*). As we detected recombination between the Stx-PP and a PP located at the *ydfJ* locus in two strains (EH1910 and 499), a different primer (the leftmost one) was used, thus labeled Type IIb. Type III analysis: In four strains, in which the PP*ompW* contained an Stx-PP, the genome of PP*ompW* and the early region of the Stx-PP were amplified using 4 primer pairs, and only these genomic regions were sequenced. Type IV and V analyses: The genomes of PP*yecE*s were amplified using 2 or 3 primer pairs to obtain the PCR products for genomic sequence determination. When the PP*yecE* contained the *stx* gene (Type IV), two *stx*-targeting primers were used as in Type II analysis. For the PP*yecE* in strain H27V05 (Type Va), only the early region was amplified and sequenced.

**S1 Table. *E. coli* strains containing the *att*-in-PP*ompW* sequence at non-*yecE* loci**.

**S2 Table. *E. coli* strains containing the *att*-in-PP*ompW* sequence at non-*yecE* loci**.

**S3 Table. *E. coli* O145:H28 strains analyzed in this study**.

**S4 Table. Complete *E. coli* genomes downloaded from the NCBI database**.

**S5 Table. Primers used for PCR amplification for prophage regions**.

**S6 Table. Primers used for long PCR analysis**.

